# Detection of airborne *Coccidioides* spores using lightweight portable air samplers affixed to uncrewed aircraft systems in California’s Central Valley

**DOI:** 10.1101/2024.10.28.620741

**Authors:** Molly T. Radosevich, Sarah Dobson, Amanda K. Weaver, Phinehas Lampman, Daniel Kollath, Lisa Couper, Grace Campbell, John W. Taylor, Justin V. Remais, Leda Kobziar, James Markwiese, Jennifer R. Head

**Author notes:** Corresponding author: Jennifer R. Head, 1415 Washington Heights, Ann Arbor, MI 48109.

## Abstract

Coccidioidomycosis is an emerging fungal infection caused by inhalation of *Coccidioides* spp. spores. While airborne dispersal is critical to *Coccidioides* transmission, limited recovery of the pathogen from air has hindered understanding of the aerosolization and transport of spores. Here, we examine uncrewed aircraft systems (UAS) with portable, active air samplers as a novel means of capturing aerosolized *Coccidioides* and characterizing emissions and exposure risk. We sampled in September 2023 in eastern San Luis Obispo County, California, in an area with confirmed *Coccidioides immitis* in soils. We completed 41 20-minute flights across 14 sites using UAS equipped with an 8 L/min bioaerosol sampler and a low-cost particulate matter sensor. We sampled source soils and air under ambient conditions using one UAS at 1-10 m above ground level, and under a simulated high-dust event using two UAS, one at <2 m height and one at 5-12 m. We detected *Coccidioides* DNA in two of 41 air samples (4.9%), both under ambient conditions at 8 m above ground level, representing the highest known height of airborne *Coccidioides* detection. Spatially explicit UAS-based sampling could enhance understanding of *Coccidioides* aerobiology and enable detection in hard-to-reach or hazardous air masses, including dust storms and wildfire smoke.

**Synopsis:** UAS-based air sampling for bioaerosols, including pathogenic *Coccidioides* spp., opens new possibilities for characterizing the aerosolization and transport of fungal spores and other airborne pathogens.

## INTRODUCTION

Coccidioidomycosis (Valley fever) is an emerging fungal infection caused by the inhalation of aerosolized spores of the *Coccidioides* genus. Infection primarily causes respiratory symptoms, but in 1-5% of cases, the disease progresses to a chronic or severe, disseminated state ^1^. The fungus is predominantly found in desert and semi-arid soils of the southwestern United States, particularly Arizona and California, where *C. posadasii* and *C. immitis*, respectively, occur ^1,2^. Since 2000, coccidioidomycosis incidence has increased nine-fold in California, most rapidly in geographic areas outside the traditionally considered endemic zones ^3,4^. This emergence appears linked to hydroclimate fluctuations and warming that promote *Coccidioides’* growth, aerosolization, and dispersal ^4–6^. Research finding positive associations between fine mineral dust exposure and coccidioidomycosis incidence ^7,8^, coupled with coccidioidomycosis outbreaks among outdoor workers exposed to dust via construction and wildfire fuel hazard reduction and fireline construction ^9,10^ suggest that soil disturbance by both natural and anthropogenic mechanisms may aerosolize fungal spores ^2^. However, there are gaps in our understanding of the conditions facilitating aerosolization and transport of *Coccidioides* under ambient air conditions and during specific events that elevate particulate matter and dust concentrations.

Understanding how, where, and when fungal spores are aerosolized and transported is crucial for developing strategies to predict and reduce exposure to *Coccidioides*. While reliable methods for detecting *Coccidioides* in soil are well established ^11^, detection of airborne fungal spores is technically challenging ^12–15^. Most studies conducting air sampling for *Coccidioides*, including large-scale surveillance efforts in Arizona ^13,16^, have relied on active air filtration using fixed-location high-volume devices that intake 100-425 L/min of air for up to 24 hours ^12,13,16,17^. While such studies have enabled valuable preliminary understanding of *Coccidioides*’ aerobiology, technology gaps and implementation costs limit the feasibility of scalable stationary, high-volume sampling networks ^18^. Meanwhile, stationary low-volume filtration and passive sampling techniques have been limited in their successful recovery of *Coccidioides* DNA ^12,14,15,19^. Thus, the processes facilitating aerosolization and transport of *Coccidioides* spp. remain poorly understood, underscoring the need for new approaches capable of capturing airborne spores.

Lightweight portable samplers show promise for collection of bioaerosols using air flow rates more relevant to human exposure risk ^20,21^. Attachment of portable samplers to the fuselage of uncrewed aircraft systems (UAS) adds flexibility to where air can be sampled, and has been used to assess the composition of microbes in smoke to understand how wildland fires mobilize organic and inorganic particles including dust and bioaerosols ^20,22,23^. However, it is currently unknown whether similar methodologies might enable detection of airborne *Coccidioides*. In this study, we assessed the feasibility of UAS equipped with lightweight, active aerosol samplers in recovering *C. immitis* from the air—a sampling approach that expands opportunities for characterizing the aerosolization and transport of pathogenic bioaerosols to and from environmental reservoirs ^23^.

## MATERIALS AND METHODS

### Site description

We conducted our study from September 22-24, 2023, in the Carrizo Plain National Monument in San Luis Obispo County, California, USA (Figure 1A), a preserved arid grassland ecosystem that provides habitat for burrowing mammal species known to be susceptible to *Coccidioides* infection ^24,25^. The Carrizo Plain is highly endemic for *Coccidioides immitis*, as previous work has isolated *C. immitis* DNA from 26% of small mammal burrow soils within the monument ^26^.

**Figure 1:**
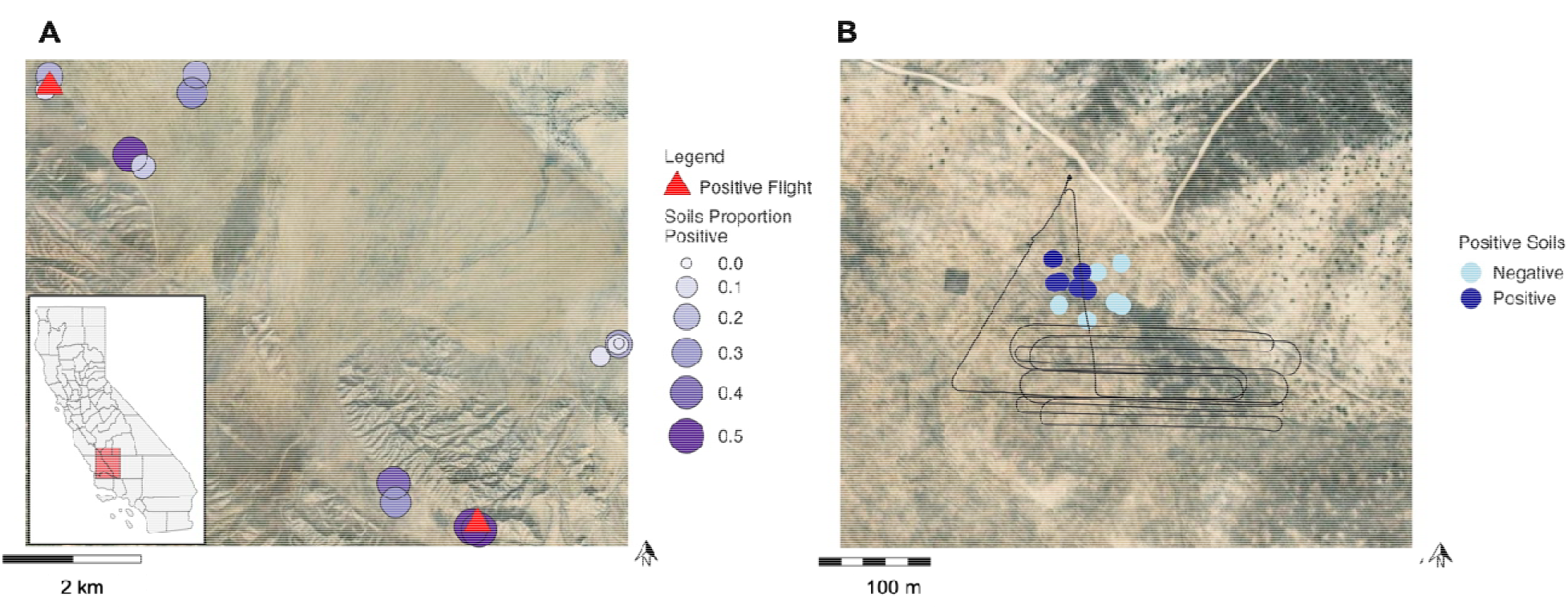
Maps of sampling locations in the Carrizo Plain, CA. (A) Layout of all sites sampled, where circles denote the proportion of positive soil samples at each site and red triangles indicate the two sites with positive flights. The inset highlights the approximate location of the Carrizo Plain National Monument in California. (B) The UAS flight path for site 14 is designated in a black line. Circles denote where soil samples were concurrently collected and whether each sample was negative (light blue) or positive (dark blue) for *Coccidioides*. Basemap source: Esri, DigitalGlobe, GeoEye, i-cubed, USDA FSA, USGS, AEX, Getmapping, Aerogrid, IGN, IGP, swisstopo, and the GIS User Community.

### Air sampling

We completed 41 flights across 14 sites using two DJI Matrice 300RTK (SZ DJI Technology Co., Ltd.) quadcopter UAS with 1.5 kg payloads, following methods described by Kobziar and colleagues ^20,22^. At each site, we implemented two sampling schemes designed to compare ambient with wind-affected air masses (Figure 2A). First, we sampled air under ambient conditions by conducting solo UAS flights at 1.2 to 10.5 m altitude. Next, we sampled air using two UAS under a paired sampling scheme designed to simulate high-dust conditions. We flew one UAS below five m altitude to mobilize dust via air turbulence from rotor downwash, and concurrently flew a second drone 5-30 m downwind at 5-10 m altitude to capture aerosolized dust potentially containing spores. Flight records were downloaded and de-encrypted using AirData software ^27^.

**Figure 2:**
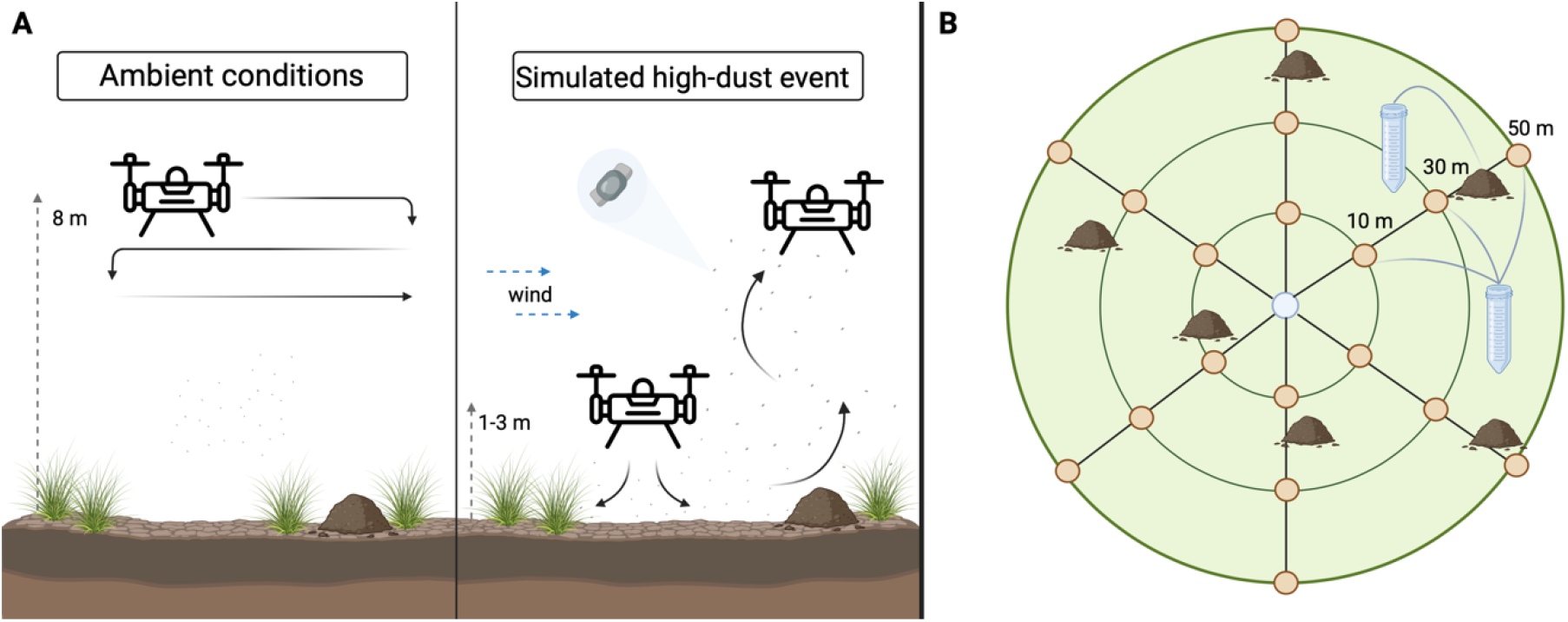
Schematics for air and soil sampling designs (A) Air sampling design for ambient and simulated high-dust flight types. (B) Systematic radial and interrupted belt transect soil sampling design. Along each transect, soil was collected from the mouths of burrows in one active precinct, and collected and pooled from the surface (top 1 cm) of soils at 10 m, 30 m, and 50 m distance from the center. Figures created using BioRender.com.

Each UAS was equipped with a battery-powered compensating Leland Legacy sampling pump and button aerosol sampler (SKC Inc., Eighty Four, PA, USA) ^22,28^ that drew in air at a volumetric rate of 8 liters per minute (L/min), split and filtered through two 2□µm PTFE (Teflon®) filters. Flights sampled 160-216 L of air over 20 to 27 minutes. A delay in air sampling was programmed to ensure that air was sampled only after the UAS reached the desired flight altitude. After each flight, both PTFE filters were brought into a portable, sterile glovebox, extracted from the button sampler, and pooled together in a 50-mL centrifuge tube before being transferred to coolers and stored on ice. The sampler was cleaned with 70% ethanol between filter loading to minimize cross-sample contamination. Samples were transferred daily to -20°C and maintained at this temperature until transferred to -80°C within four days. To help account for possible contamination, two to three procedural blanks were collected daily in the field. Briefly, new filters were installed in the clean button sampler and allowed to sit for 1 minute, then removed into 50-mL centrifuge tubes in a sterilized glovebox using the same procedures as for positive samples. Blanks and samples were stored and analyzed identically.

### Measurement of environmental covariates

During flight, the UAS recorded continuous measures of speed, latitude, longitude, and height above ground level. Each UAS had an attached PurpleAir PA-II-SD (PurpleAir, Inc.), which included two PMS5003 particulate matter (PM) optical sensors (Plantower, Beijing, China) ^22,28^. The PurpleAir recorded data every eight seconds on air temperature, relative humidity, dewpoint, and particulate matter smaller than 1.0, 2.5, and 10 microns (PM_1.0_, PM_2.5_, and PM_10.0_). Technical issues prevented one PurpleAir monitor from recording data during eight of the 41 flights. All UAS data manipulation and analysis was done in Python (version 3.10) and R (version 4.3.2). We acquired hourly average and maximum wind speed from the Remote Automatic Weather Stations (RAWS) US Climate Archive, which has a station located within the Carrizo Plain ^29^. These observations were linked to the UAS flights based on closest match to time of flight.

### Soil sampling

To examine whether *C. immitis* in air was associated with *C. immitis* in soils, we collected soil samples (12 per site, total n=168) at locations beneath the UAS airspace of each flight site. We prioritized sampling soil associated with rodent burrows, where prior work has found a high probability of detecting *Coccidioides* ^26,30,31^. Per site, we collected six samples from inside burrow entrances (prioritizing burrows that appeared active via presence of feet and tail markings, clipped vegetation, and fecal pellets) and six from nearby surface soil (prioritizing loose soils that could be readily dispersed and appeared to have been excavated from burrows). Our sampling design followed a combined radial and interrupted belt transect approach (Figure 2B) adapted from prior work ^32^. Herein, six 50-m radii were measured from a centroid at 60-degree angles. Along each transect, we collected and pooled soil from the top one cm of the surface at approximately 10 m, 30 m, and 50 m (total volume 25-35 mL). We also sampled one rodent precinct (i.e., cluster of burrows) per transect, pooling samples from up to three burrow entrances per precinct (total volume 10-30 mL). Soils were collected using a stainless-steel scoop with a 10-inch handle, placed into sterile, 50-mL centrifuge tubes, and sealed in zip-top plastic bags. Scoops were thoroughly cleaned with 70% ethanol between samples to prevent contamination. Samples were stored at ambient conditions until analysis, following standard protocols for the analysis of *Coccidioides* in soils ^32–34^.

### DNA extraction and detection of *Coccidioides DNA*

Genomic DNA was extracted from soils using the DNeasy PowerSoil Pro Kit (QIAGEN, Hilden, Germany), and from filters using a phenol-chloroform extraction procedure ^35^ (personal correspondence: Bridget Barker and Daniel Kollath). A laboratory blank was processed alongside the samples and field blanks. Tubes containing the filters were washed with lysis buffer to maximize DNA yield. DNA was quantified using a Qubit™ fluorometer (ThermoFisher Scientific, Waltham, Massachusetts, USA). Samples were diluted to 12 ng/µL and assessed for presence of *Coccidioides* DNA using the CocciEnv qPCR assay ^11^. Samples were run in quadruplicate and classified as positive for *Coccidioides* if at least three of four replicates had a cycle threshold (Ct) value below 40, following prior work ^14^.

### Statistical analyses

We mapped flight paths and summarized flight characteristics, environmental data, and sample positivity rates across sites. We compared the prevalence of *C. immitis* across sample types and examined the relationships between environmental factors (PM concentration, relative humidity, temperature, height above ground, time of day, and wind speeds), flight characteristics (altitude, flight duration, distance travelled, and volume of air sampled) and detection of *C. immitis* in air samples using univariate logistic regression. Missing data resulting from the UAS PurpleAir monitor operating error were removed from analysis. The US Environmental Protection Agency’s correction equations were used to estimate PM_2.5_ concentrations from PurpleAir sensor particulate data ^36^. Statistical analysis was performed in R (version 4.3.2) ^37^.

## RESULTS

Out of 41 air samples collected via UAS, two were positive for *C. immitis* DNA (4.9%) (Table 1). Both positive samples were collected during solo ambient flights at mean heights of 7.9 m and 8.4 m above ground, at two different sites (Figure 1). Meanwhile, 45 out of 168 soil samples were positive for *Coccidioides* DNA (27%) (Table 1). The proportion of positive soil samples at any given site ranged from 0-66% (Figure 1). At both sites from which positive air samples were collected, *C. immitis* DNA was detected in the soil beneath the UAS airspace. All laboratory and field blanks collected had no to negligible DNA quantities (<0.07 ng/uL) and all were negative for *Coccidioides* DNA (see Methods).

**Table 1:**
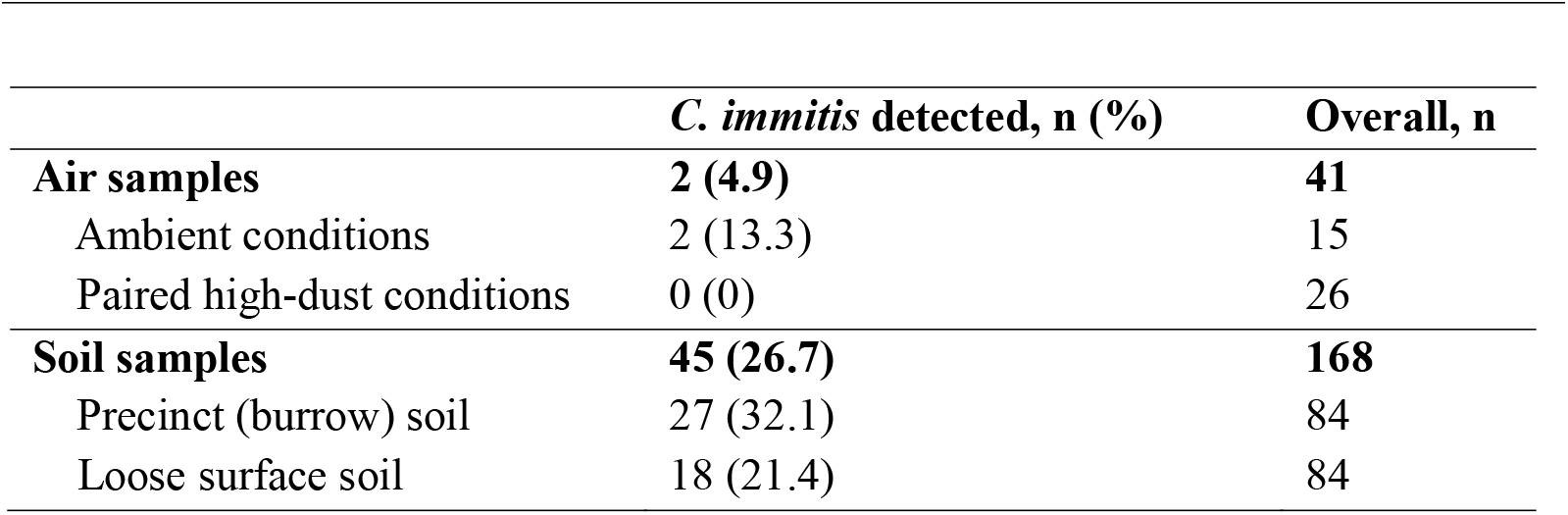
Percentage of air and soil samples collected in southeastern San Luis Obispo County, California, September 2023 with *C. immitis* DNA.

We found no significant associations between *Coccidioides* detection in airborne samples and environmental data or flight characteristics, although we lacked statistical power to do so. Mean flight duration was 22.4 minutes (range: 20-27 minutes), corresponding to 160-216 L of filtered air. The mean height above ground for ambient flights was 6.3 m (range: 1.2-10.5 m). For the paired sampling scheme, the mean height above ground for low flights was 1.08 (range: 0.84-1.44 m) and for high flights was 7.53 (range: 5.21-11.77 m). The mean concentrations of PM_1.0,_ PM_2.5_, and PM_10.0_ were 21.8 +/-4.4 µg/m^3^, 21.3 +/-3.0 µg/m^3^, and 40.0 +/-7.1 µg/m^3^, respectively. There were no significant differences in mean PM_1.0,_ PM_2.5_, or PM_10.0_ concentration based on flight type for either drone. Using data gathered from the Carrizo RAWS weather station, the average hourly temperature during sampling was 25.3°C, with 31.4% relative humidity, and average wind speeds of 7.3 mph.

## DISCUSSION

This is the first study, to our knowledge, to detect *C. immitis* in the air using portable active air samplers attached to UAS. We recovered *C. immitis* DNA from two of 41 air samples collected by a UAS-based sampler, both collected under ambient conditions at heights of approximately 8 m above ground – the highest height at which airborne *Coccidioides* has been detected. Prior surveillance efforts have sampled air using fixed devices stationed at 1.4 m and 4.6 m above the ground ^13,16^. This study establishes UAS as a feasible sampling method for *Coccidioides*, laying groundwork for future investigations to evaluate the atmospheric and environmental factors influencing emissions and transport of spores.

Contrary to our expectations, *C. immitis* was only detected during ambient flights and not during flights conducted specifically to disturb and aerosolize soils presumably containing spores. We did not observe any difference in PM concentrations between ambient and lower-height flights, suggesting that the relevant size fraction of dust was not aerosolized during paired flights. We found no association between soil positivity and airborne presence of *Coccidioides*, though this may be due to limited power in having only two positive air samples. However, given the potential for long-distance dispersal of fungal spores in the air ^38^, the relatively high heights of detection, and evidence that the air mycobiome has a distinct composition from the soil mycobiome under ambient conditions ^19^, it is possible that the spores we detected were dispersed via wind from farther distances.

We propose that use of UAS in air sampling can help understand environmental factors affecting *Coccidioides* dispersal in endemic regions, given its spatial precision via integrated GPS tracking functionality; ability to sample at varying heights above ground level and hard-to-reach locations; and ability to concurrently collect data from monitors affixed to the UAS ^22^. One compelling application of this technology could be to assess the role of wildfires and prescribed burns in dispersing fungal spores and other bioaerosols, a key knowledge gap that has only begun to be addressed ^23,39^. In the process of burning organic soils and plant material, fires may directly emit fungal matter and draw matter into the smoke column via convective winds. Such bioaerosols have the potential to travel hundreds of miles on wind currents before being deposited or inhaled ^20,40–42^. Past work has demonstrated the efficacy of UAS in sampling viable microbiological aerosols within smoke plumes ^20,22^, finding that plumes contain higher concentrations and a greater diversity of microorganisms than ambient air ^22^. Employing UAS to quantify the potential concentrations of aerosolized *Coccidioides* during wildfires would enable practical linkages with smoke models parameterized for pathogenic bioaerosol transport ^43^.

Our findings also suggest that portable button aerosol samplers utilizing lower flow rates (even without UAS integration) could be leveraged to assess personal exposure to *Coccidioides* spores. We demonstrate isolation of *C. immitis* in only 176-184 L of air using lightweight portable samplers that filter air at a rate akin to human breathing (8 L/min). Our study found *C. immitis* DNA in 4.9% of air samples, similar to the positivity rate in Phoenix, AZ (7.4%), which relied on high-volume fixed-location samplers that filtered 2,400 liters of air ^16^.

While we demonstrated a novel approach for detecting airborne *Coccidioides*, our small sample size precluded us from detecting associations between *Coccidioides* presence and environmental factors. Our findings inform compelling areas of future research, including investigations of the relationships between airborne *Coccidioides* presence and various land use types, environmental factors, dust-generating events such as wildfires, and presence in soil. Given the public health threat of the emergence of coccidioidomycosis in the western United States, it is imperative to continue investigating how environmental and anthropogenic disturbances may facilitate the transport of *Coccidioides* spores and influence exposure risk.

## ACKNOWLEDGEMENTS

This research was supported in part by the US Environmental Protection Agency Ecology Effects Branch, Pacific Ecological Systems Division (Corvallis, OR) and the National Institute of Allergy and Infectious Diseases of the National Institutes of Health under award no. R01AI148336. JRH was funded by NIH Award K01AI173529. Additional support was provided by the Keck Foundation. UAS pilots included Tim Wallace of Black Mountain UAS, LLC. Figure 2 and the Graphical Abstract were created using BioRender.com. Maps in Figure 1 were created using ArcGIS® software by Esri. ArcGIS® and ArcMap™ are the intellectual property of Esri and are used herein under license. Copyright © Esri. All rights reserved. For more information about Esri® software, please visit www.esri.com. The views expressed in this article are those of the authors and do not necessarily represent the views or policies of the U.S. Environmental Protection Agency.

## References

1. Valley Fever (Coccidioidomycosis) | Types of Fungal Diseases | Fungal | CDC. February 3, 2021. Accessed February 12, 2024. https://www.cdc.gov/fungal/diseases/coccidioidomycosis/index.html

2. Galgiani JN, Ampel NM, Blair JE, et al. Coccidioidomycosis. Clin Infect Dis. 2005;41(9):1217–1223. doi:10.1086/496991

3. Cooksey GLS. Regional Analysis of Coccidioidomycosis Incidence — California, 2000– 2018. MMWR Morb Mortal Wkly Rep. 2020;69. doi:10.15585/mmwr.mm6948a4

4. Gorris ME, Treseder KK, Zender CS, Randerson JT. Expansion of Coccidioidomycosis Endemic Regions in the United States in Response to Climate Change. GeoHealth. 2019;3(10):308–327. doi:10.1029/2019GH000209

5. Head JR, Sondermeyer-Cooksey G, Heaney AK, et al. Effects of precipitation, heat, and drought on incidence and expansion of coccidioidomycosis in western USA: a longitudinal surveillance study. Lancet Planet Health. 2022;6(10):e793–e803. doi:10.1016/S2542-5196(22)00202-9

6. Weaver EA, Kolivras KN. Investigating the Relationship Between Climate and Valley Fever (Coccidioidomycosis). EcoHealth. 2018;15(4):840–852. doi:10.1007/s10393-018-1375-9

7. Colson AJ, Vredenburgh L, Guevara RE, Rangel NP, Kloock CT, Lauer A. Large-Scale Land Development, Fugitive Dust, and Increased Coccidioidomycosis Incidence in the Antelope Valley of California, 1999–2014. Mycopathologia. 2017;182(5-6):439–458. doi:10.1007/s11046-016-0105-5

8. Tong DQ, Gorris ME, Gill TE, Ardon□Dryer K, Wang J, Ren L. Dust Storms, Valley Fever, and Public Awareness. GeoHealth. 2022;6(8):e2022GH000642. doi:10.1029/2022GH000642

9. Laws RL, Jain S, Cooksey GS, et al. Coccidioidomycosis outbreak among inmate wildland firefighters: California, 2017. Am J Ind Med. 2021;64(4):266–273. doi:10.1002/ajim.23218

10. Freedman M, Jackson BR, McCotter O, Benedict K. Coccidioidomycosis Outbreaks, United States and Worldwide, 1940–2015. Emerg Infect Dis. 2018;24(3):417–424. doi:10.3201/eid2403.170623

11. Bowers JR, Parise KL, Kelley EJ, et al. Direct detection of Coccidioides from Arizona soils using CocciENV, a highly sensitive and specific real-time PCR assay. Med Mycol. 2019;57(2):246–255. doi:10.1093/mmy/myy007

12. Chow NA, Griffin DW, Barker BM, Loparev VN, Litvintseva AP. Molecular detection of airborne Coccidioides in Tucson, Arizona. Med Mycol. 2016;54(6):584–592. doi:10.1093/mmy/myw022

13. Gade L, McCotter OZ, Bowers JR, et al. The detection of Coccidioides from ambient air in Phoenix, Arizona: Evidence of uneven distribution and seasonality. Med Mycol. 2020;58(4):552–559. doi:10.1093/mmy/myz093

14. Wagner R, Montoya L, Head JR, Campo S, Remais J, Taylor JW. Coccidioides undetected in soils from agricultural land and uncorrelated with time or the greater soil fungal community on undeveloped land. PLOS Pathog. 2023;19(5):e1011391. doi:10.1371/journal.ppat.1011391

15. Ajello L, Maddy K, Crecelius G, Hugenholtz PG, Hall LB. Recovery of Coccidioides immitis from the air. Sabouraudia. 1965;4(2):92–95. doi:10.1080/00362176685190231

16. Porter WT, Gade L, Montfort P, et al. Understanding the exposure risk of aerosolized Coccidioides in a Valley fever endemic metropolis. Sci Rep. 2024;14(1):1311. doi:10.1038/s41598-024-51407-x

17. Daniels JI, Wilson WJ, DeSantis TZ, et al. Development of a Quantitative TaqMan{trademark}-PCR Assay and Feasibility of Atmosphoric Collection for Coccidioides Immits for Ecological Studies.; 2002:UCRL-ID-146977, 15002759. doi:10.2172/15002759

18. DHS Exploring New Methods to Replace BioWatch and Could Benefit from Additional Guidance. Published online May 2021. https://www.gao.gov/products/gao-21-292

19. Wagner R, Montoya L, Gao C, Head JR, Remais J, Taylor JW. The air mycobiome is decoupled from the soil mycobiome in the California San Joaquin Valley. Mol Ecol. 2022;31(19):4962–4978. doi:10.1111/mec.16640

20. Kobziar LN, Vuono D, Moore R, et al. Wildland fire smoke alters the composition, diversity, and potential atmospheric function of microbial life in the aerobiome. ISME Commun. 2022;2(1):1–9. doi:10.1038/s43705-022-00089-5

21. Yao M, Mainelis G. Use of portable microbial samplers for estimating inhalation exposure to viable biological agents. J Expo Sci Environ Epidemiol. 2007;17(1):31–38. doi:10.1038/sj.jes.7500517

22. Kobziar LN, Pingree MRA, Watts AC, Nelson KN, Dreaden TJ, Ridout M. Accessing the Life in Smoke: A New Application of Unmanned Aircraft Systems (UAS) to Sample Wildland Fire Bioaerosol Emissions and Their Environment. Fire. 2019;2(4):56. doi:10.3390/fire2040056

23. Bonfantine K, Vuono DC, Christner BC, et al. Evidence for Wildland Fire Smoke Transport of Microbes From Terrestrial Sources to the Atmosphere and Back. J Geophys Res Biogeosciences. 2024;129(9):e2024JG008236. doi:10.1029/2024JG008236

24. Endicott R, Dillard D, Barnes M, et al. Carrizo Plain Ecosystem Project. Published online 2017.

25. Emmons CW. Coccidioidomycosis in Wild Rodents. A Method of Determining the Extent of Endemic Areas. Public Health Rep 1896-1970. 1943;58(1):1–5. doi:10.2307/4584326

26. Head JR, Camponuri SK, Weaver AK, et al. Small mammals and their burrows shape the distribution of Coccidioides in soils: a long-term ecological experiment. Published online September 24, 2024. doi:10.1101/2024.09.21.613892

27. Drone Data Management and Flight Analysis | Airdata UAV. Accessed September 15, 2024. https://airdata.com/

28. Nelson KN, Boehmler JM, Khlystov AY, et al. A Multipollutant Smoke Emissions Sensing and Sampling Instrument Package for Unmanned Aircraft Systems: Development and Testing. Fire. 2019;2(2):32. doi:10.3390/fire2020032

29. RAWS USA Climate Archive State Selection Map. Accessed April 1, 2024. https://raws.dri.edu/

30. del Rocío Reyes-Montes M, Pérez-Huitrón MA, Ocaña-Monroy JL, et al. The habitat of Coccidioides spp. and the role of animals as reservoirs and disseminators in nature. BMC Infect Dis. 2016;16(1):550. doi:10.1186/s12879-016-1902-7

31. Barker BM, Tabor JA, Shubitz LF, Perrill R, Orbach MJ. Detection and phylogenetic analysis of Coccidioides posadasii in Arizona soil samples. Fungal Ecol. 2012;5(2):163–176. doi:10.1016/j.funeco.2011.07.010

32. Chow NA, Kangiser D, Gade L, et al. Factors Influencing Distribution of Coccidioides immitis in Soil, Washington State, 2016. mSphere. 2021;6(6):e00598–21. doi:10.1128/mSphere.00598-21

33. Friedman L, Smith CE, Pappagianis D, Berman RJ. Survival of Coccidioides immitis Under Controlled Conditions of Temperature and Humidity. Am J Public Health Nations Health. 1956;46(10):1317–1324. doi:10.2105/AJPH.46.10.1317

34. Greene DR, Koenig G, Fisher MC, Taylor JW. Soil isolation and molecular identification of Coccidioides immitis. Mycologia. 2000;92(3):406–410. doi:10.1080/00275514.2000.12061175

35. Mead HL, Blackmon AV, Vogler AJ, Barker BM. Heat Inactivation of Coccidioides posadasii and Coccidioides immitis for Use in Lower Biosafety Containment. Appl Biosaf J Am Biol Saf Assoc. 2019;24(3):123. doi:10.1177/1535676019856525

36. US EPA O. AirNow Fire and Smoke Map: Extension of the U.S.-Wide Correction for PurpleAir PM2.5 Sensors Webinar Archive. May 25, 2021. Accessed April 1, 2024. https://www.epa.gov/research-states/airnow-fire-and-smoke-map-extension-us-wide-correction-purpleair-pm25-sensors

37. R: The R Project for Statistical Computing. Accessed April 25, 2024. https://www.r-project.org/

38. Golan JJ, Pringle A. Long-Distance Dispersal of Fungi. Heitman J, Crous PW, eds. Microbiol Spectr. 2017;5(4):5.4.03. doi:10.1128/microbiolspec.FUNK-0047-2016

39. Ellington AJ, Walters K, Christner BC, et al. Dispersal of microbes from grassland fire smoke to soils. ISME J. Published online October 15, 2024:wrae203. doi:10.1093/ismejo/wrae203

40. Kobziar LN, Thompson GR. Wildfire smoke, a potential infectious agent. Science. 2020;370(6523):1408–1410. doi:10.1126/science.abe8116

41. Moore RA, Bomar C, Kobziar LN, Christner BC. Wildland fire as an atmospheric source of viable microbial aerosols and biological ice nucleating particles. ISME J. 2021;15(2):461–472. doi:10.1038/s41396-020-00788-8

42. Cottle P, Strawbridge K, McKendry I. Long-range transport of Siberian wildfire smoke to British Columbia: Lidar observations and air quality impacts. Atmos Environ. 2014;90:71–77. doi:10.1016/j.atmosenv.2014.03.005

43. Kobziar LN, Lampman P, Tohidi A, et al. Bacterial Emission Factors: A Foundation for the Terrestrial-Atmospheric Modeling of Bacteria Aerosolized by Wildland Fires. Environ Sci Technol. Published online January 24, 2024:acs.est.3c05142. doi:10.1021/acs.est.3c05142

